# Anti-prion drugs do not improve survival in knock-in models of inherited prion disease

**DOI:** 10.1101/2023.09.28.559951

**Authors:** Daniel J. Walsh, Judy R. Rees, Surabhi Mehra, Matthew E.C. Bourkas, Lech Kaczmarczyk, Erica Stuart, Walker S. Jackson, Joel C. Watts, Surachai Supattapone

**Affiliations:** Department of Biochemistry Geisel School of Medicine at Dartmouth, Hanover, New Hampshire 03755, USA; Department of Epidemiology Geisel School of Medicine at Dartmouth, Hanover, New Hampshire 03755, USA; Department of Community and Family Medicine Geisel School of Medicine at Dartmouth, Hanover, New Hampshire 03755, USA; Tanz Centre for Research in Neurodegenerative Diseases, University of Toronto, Toronto, Ontario, Canada; Department of Biochemistry, University of Toronto, Toronto, Ontario, Canada; Wallenberg Center for Molecular Medicine, Department of Biomedical and Clinical Sciences, Linköping University Sweden; Department of Medicine, Geisel School of Medicine at Dartmouth, Hanover, New Hampshire 03755, USA

**Keywords:** prion, genetic, mutant, drug, therapy, familial CJD, fatal familial insomnia

## Abstract

Prion diseases uniquely manifest in three distinct forms: inherited, sporadic, and infectious. Wild-type prions are responsible for the sporadic and infectious versions, while mutant prions cause inherited variants like fatal familial insomnia (FFI) and familial Creutzfeldt-Jakob disease (fCJD). Although some drugs can prolong prion incubation times up to four-fold in rodent models of infectious prion diseases, no effective treatments for FFI and fCJD have been found.

In this study, we evaluated the efficacy of various anti-prion drugs on newly-developed knock-in mouse models for FFI and fCJD. These models express bank vole prion protein (PrP) with the pathogenic D178N and E200K mutations. We applied various drug regimens known to be highly effective against wild-type prions *in vivo* as well as a brain-penetrant compound that inhibits mutant PrP^Sc^ propagation *in vitro*. None of the regimens tested (Anle138b, IND24, Anle138b + IND24, cellulose ether, and PSCMA) significantly extended disease-free survival or prevented mutant PrP^Sc^ accumulation in either knock-in mouse model, despite their ability to induce strain adaptation of mutant prions. Paradoxically, the combination of Anle138b and IND24 appeared to accelerate disease by 16% and 26% in kiBVI^E200K^ and kiBVI^D178N^ mice, respectively, and accelerated the aggregation of mutant PrP molecules *in vitro*. Our results show that anti-prion drugs originally developed to treat infectious prion diseases do not necessarily work for inherited prion diseases, and that the recombinant sPMCA is not a reliable platform for identifying compounds that target mutant prions. This work underscores the need to develop therapies and validate screening assays specifically for mutant prions.

## Introduction

Prion diseases are fatal neurodegenerative diseases that uniquely occur in inherited, sporadic, and infectious forms(1). All forms of prion diseases are caused by autocatalytic misfolding of the prion protein (PrP), a host-encoded glycoprotein. Inherited prion diseases such as familial Creutzfeldt-Jakob disease (fCJD), fatal familial insomnia (FFI), and Gerstmann-Sträussler-Scheinker disease (GSS) are all caused by pathogenic PrP mutations(2–4), whereas sporadic and infectious forms of prion disease are caused by the misfolding of normal wild-type PrP (PrP^C^) into an infectious conformer (PrP^Sc^)(5, 6). Interestingly, mutant prions from patients with inherited prion diseases can also be infectious; both FFI and fCJD prions have been experimentally transmitted to normal hosts(7, 8).

Several easily administered, non-toxic drug treatments have been shown to prolong lifespan in mice inoculated with scrapie, an infectious form of prion disease. For instance, IND24, a 2-aminothiazole compound, and Anle138b, a 3,5-diphenyl-pyrazole (DPP) derivative, both extend scrapie incubation times >2-fold when administered orally immediately after prion inoculation(9) (10, 11); and prophylactic subcutaneous administration of polymeric cellulose ether extends scrapie incubation times 4-fold(12). Despite these advances, there are inherent difficulties in treating infectious and sporadic forms of prion diseases. First, wild type PrP^Sc^ molecules can adapt into drug-resistant conformations during treatment, limiting drug efficacy(10). Remarkably, even combination and alternating chemotherapy regimens cannot prevent the emergence of drug-resistant PrP^Sc^ molecules(11, 13). Second, patients with sporadic and infectious forms of prion disease typically present with neurological symptoms at late clinical stages when the majority of PrP^Sc^ accumulation and irreversible neurodegeneration has already occurred.

On the other hand, inherited prion diseases represent a more attractive target for therapeutic intervention. First, patients with inherited prion disease are usually diagnosed by genetic testing many years before the onset of clinical symptoms, and therefore treatment for inherited prion disease can be initiated much earlier than for sporadic or infectious forms of prion diseases. Although experiments using genetically engineered CRE-Lox mice have shown that, in principle, the process of neuronal dysfunction during the symptomatic phase of prion disease is reversible if PrP^Sc^ production can be completely halted(14), other studies indicate that anti-prion drug therapy is typically more effective if administered before the onset of clinical symptoms(15). For instance, Giles et al. reported that prophylactic administration of IND24 to prion-infected animals extended lifespan ∼4-fold, whereas administration of IND24 1 day after inoculation extended lifespan ∼1.7-fold(16). Second, it is possible that mutant prions are less malleable than wild-type prions, based on the following observations: (1) each pathogenic PrP mutation appears to cause a characteristic strain phenotype and mutant PrP^Sc^ conformation *in vivo*(17–21), as discussed above (and notably even E200G produces a different phenotype than E200K(22)); (2) mutations and polymorphisms have been shown to distort the folding landscape of PrP molecules, thermodynamically favoring conversion into a sequence-specific misfolded conformation(23); and (3) unlike wild-type prions, mutant prions can be propagated *in vitro* without cofactor molecules(24), and therefore may not be able to harness cofactor diversity to create alternative conformers. If mutant prions are truly not malleable, it is possible that inherited prion diseases could be indefinitely suppressed by static drug therapy without strain adaptation or the emergence of drug-resistant strains.

Although treatment of inherited prion diseases appears to be a tractable goal, there have been no drugs specifically developed to target mutant prions. Recently, Vallabh *et al.* accurately noted that a longstanding barrier to developing and testing such drugs pre-clinically has been the lack of animal models of inherited prion disease that exhibit shortened lifespans without transgene overexpression(25). Here, we evaluate several drug regimens that can either inhibit wild-type PrP^Sc^ accumulation and prolong incubation time in scrapie-infected animals or inhibit mutant PrP^Sc^ propagation in sPMCA reactions for their ability to treat new knock-in mouse models of FFI and fCJD with shortened lifespans.

## Materials and Methods

### Ethical statement

The Guide for the Care and Use of Laboratory Animals of the National Research Council was strictly followed for animal experiments. The mouse bioassay experiment in this study was conducted in accordance with protocol supa.su.1 as reviewed and approved by Dartmouth College’s Institutional Animal Care and Use Committee, operating under the regulations/guidelines of the NIH Office of Laboratory Animal Welfare (assurance number A3259-01).

### Mouse models

The methods used to design and produce kiBVI^D178N^ and kiBVI^E200K^ mice were originally described in a complementary manuscript(26). Briefly, gene targeting in V6.5 embryonic stem cells was performed at the DZNE/Bonn University using CRISPR technology as described previously(27). Plasmids containing the open reading frames of either wild-type, D178N-mutant, or E200K-mutant BVPrP (I109 polymorphic variant) were used as a starting point(28). Targeting constructs were generated by ligating the respective BVPrP open reading frame variants between EagI and ClaI sites of the intermediate vector pWJPrP101(27) containing homology regions and a neomycin selection cassette removable by Flp recombinase. The Cas9 vector used for double-strand break generation in the *Prnp* gene is available from Addgene (plasmid #78621) (27). Expansion of gene-edited embryonic stem cells and aggregation with diploid CD-1(ICR) mouse embryos was performed at The Centre for Phenogenomics (Toronto, Canada). Chimeric mice were identified by their mixed coat colors and then bred with B6(Cg)-*Tyr^c-2J^*/J mice (“B6-albino mice”; Jackson Lab #000058) to identify those with germline transmission of the gene-edited *Prnp* allele. Successful chimeras were crossed with a Flp deleter strain (B6.129S4-*Gt(ROSA)26Sor^tm1(FLP1)Dym^*/RainJ; Jackson Lab #009086) to remove their selectable marker and then backcrossed with wild-type C57BL/6 mice to remove the Flp transgene. Mice that were positive for the BVPrP knock-in allele and negative for the Flp transgene were then intercrossed to create homozygous knock-in mice. All knock-in lines were maintained by crossing homozygous female with homozygous male mice.

### Drug Treatment, Diagnosis, and Neuropathology

Female mice were used for this study and housed in groups of 4. Starting at 1 month of age, mice were fed Teklad chow (Envigo, Madison, WI) formulated by Envigo to contain either 280 mg compound/kg body weight/day IND24, 400 mg/kg/day Anle138b, or 280 mg/kg/day IND24 plus 400 mg/kg/day Anle138b. IND24 and Anle138b were synthesized by Sundia (Sacramento, CA). Control animals were fed Teklad chow alone. Mice were administered Metolose® 60SH-50 (cellulose ester viscosity 50) (Shin Etsu, Akron, OH) by subcutaneous injections (4 g/kg body weight/injection) every 4 months starting at 1 month of age. Mice were administered (poly [4-styrenesulfonic acid-co-maleic acid] (PSCMA) (Sigma Aldrich, St. Louis, MO) (∼480 mg/kg body weight/day) starting at 1 month of age through drinking water (2mg/mL).

Mice were monitored daily for the appearance and progression of neurological symptoms associated with prion disease, including ataxia, kyphosis, limb paralysis, and slow gait. Neuropathology was performed as previously described(29). Microscopic sections were commercially prepared by iHisto (Salem, MA), and vacuolation in various brain regions was scored on a 0-5 scale. Mean vacuolation scores were plotted using the *radarchart* function of the fmsb package (https://cran.r-project.org/web/packages/fmsb/) in R (version 4.1.3).

### Detection of mutant PrP^Sc^ in brain homogenates

Brains for biochemical analysis were homogenized to 10% (w/v) in sterile PBS. Sodium deoxycholate and NP-40 were each added to final concentration of 0.5% and samples were incubated for 20 min with intermittent vortexing. After a 5 min centrifugation at 1,000 x *g*, supernatants were recovered, and total protein was quantified using a Micro BCA Protein Assay Kit (ThermoFisher). The remaining supernatant was either treated with thermolysin (TL) protease (Sigma) at a ratio of 50 µg protein to 1 µg protease or water (-TL control), shaking for 1 hr at 600 rpm and 37°C. Digests were quenched by addition of 5 mM EDTA. Sarkosyl was then added to a final concentration of 2% (w/v) and samples were incubated for 5 min before centrifugation at 100,000 x *g* for 45 min. Supernatants were aspirated and pellets were resuspended in Laemmli SDS sample buffer (Bioland Scientific) and boiled for 15 min at 95°C. Unless otherwise specified, all steps above were performed on ice or at 4°C.

Denatured protein pellets were run on 12% SDS-PAGE gels and transferred to PVDF membrane using a semi-dry blotting apparatus. Western blots were probed using either EP1802Y (Abcam, 1:10,000 dilution) or 27/33 (in-house mAb, epitope = 142-149 mouse numbering, 1:25000 dilution) primary antibodies and HRP-linked goat anti-rabbit or sheep anti-mouse secondary antibodies, respectively.

### Mutant recombinant sPMCA assay

Unseeded sPMCA reactions using recombinant D177N mouse (Mo) PrP substrate were performed and analyzed as previously described(24) with the modification that the final PrP concentration in each reaction was 40 µg/mL.

### QuIC assay

Quaking-induced conversion reactions were performed as described previously (30), with the following modifications. Recombinant PrP substrates containing the D178N or E200K mutations were created by site-directed mutagenesis with the GeneTailor system (Invitrogen) using the bank vole (BV) I109 recPrP expression plasmid(30) as template. The D178N mutation was inserted using 5’-AGAACAACTTCGTGCACAATTGCGTCAACATCACC-3’ and 5’-GGTGATGTTGACGCAATTGTGCACGAAGTTGTTCT-3’ as forward and reverse primers, respectively. The E200K mutation was inserted using 5’-AGGGGGAGAACTTCACGAAGACCGACGTCAAGATG-3’ and 5’-CATCTTGACGTCGGTCTTCGTGAAGTTCTCCCCCT-3’ as forward and reverse primers, respectively. The mutant genes were cloned into the pET-22b plasmid for bacterial expression and purified, as described previously (30–32).

Reactions were allowed to misfold spontaneously by the omission of PrP^Sc^ seeds, and were supplemented with either 1 µM Anle13b, 1 µM IND24, 1 µM Anle13b + 1 µM IND24, or DMSO as a vehicle control (“Untreated”). Shaking was performed in an Allsheng MSC-100 Thermo-Shaker set at 42°C and 900 rpm for 30 hours. At 2 hr increments, Thioflavin T fluorescence was measured in a SpectraMax iD5 plate reader (Molecular Devices) pre-warmed to 42°C.

t_1/2_ was measured as the time from the start of the experiment to the point at which half-maximal fluorescence was reached. Average t_1/2_ and standard error of the mean (SEM) was calculated for each treatment condition across 3 replicate reactions.

### Statistical analyses

We used Mann-Whitney tests with statistical significance set at alpha = 0.05 for all statistical comparisons between each treatment group and untreated controls. Three types of comparisons were evaluated: (i) incubation times from inoculation to death, (ii) vacuolation scores in each region of the brain, and (iii) t_1/2_ values from QuIC assays.

## Acknowledgements

This study was funded by the National Institute for Neurological Diseases and Stroke R37NS125431, R01NS117276, and R01NS118796 to S.S.) and the National Institutes of Health and P20-GM113132 (P.I. Dean Madden). The authors thank Rachel Pepin and Emma Fiske for assistance with protease-digestion experiments, and Ciara Groesbeck and Eric DuFour for their assistance with veterinary care and tissue harvests.

## Results

For this study, we used recently developed knock-in mouse models of inherited prion diseases with shortened lifespans(26). These models express I109 bank vole (BV) PrP with either the D178N or E200K pathogenic mutations (hereafter, these models are respectively termed “kiBVI^D178N^ mice” and “kiBVI^E200K^ mice” for brevity, but is important to note the caveat that these models do not produce the same mutant PrP^Sc^ conformation and neuropathological patterns seen in human patients.) In choosing drug regimens to test in kiBVI^D178N^ and kiBVI^E200K^ mice, we included a variety of oral and sub-cutaneous treatments previously shown to be highly effective in treating prion-infected mice. These regimens (Anle138b, IND24, a combination regimen of Anle138b plus IND24, and Metolose cellulose ester) all produce survival indices(33) between 2-4-fold in scrapie-infected rodents(10–12, 34) (summarized in **Table 1, column 2**). We also included PSCMA, an oral, brain-penetrant compound previously shown to inhibit PrP^C^-dependent Aβ oligomer toxicity *in vivo*(35) because we found that it could inhibit spontaneous D177N Mo PrP^Sc^ propagation in sPMCA reactions **(Figure 1)**.

**Figure 1:**
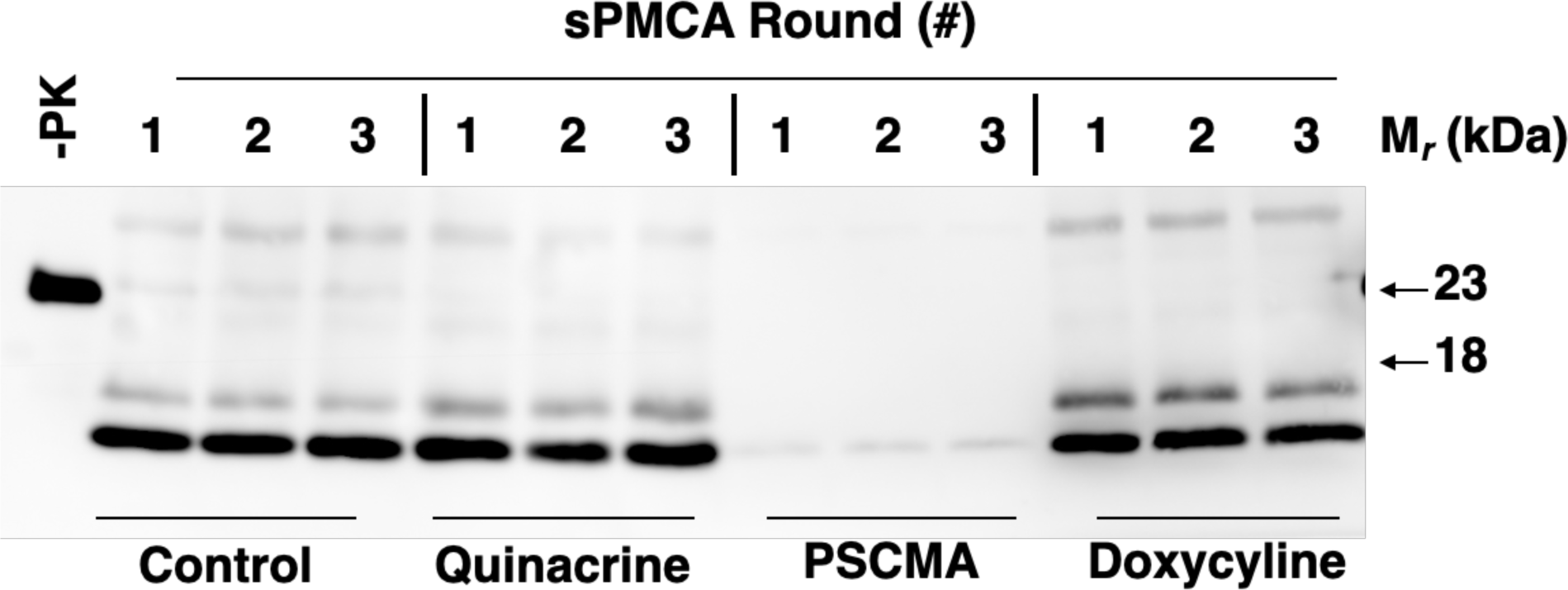
Effect of various drug treatments on mutant recombinant sPMCA propagation. Western blots of unseeded 3-round sPMCA reactions using recombinant D177N Mo PrP substrate. Where indicated, reactions contained 20 µM quinacrine, PSCMA, or doxycycline. Blot probed with anti-PrP mAb 27-33. -PK = control sample not subjected to protease digestion. All other samples treated with 25 µg/mL proteinase K.

**Table 1.** Effect of drug treatments on infectious and inherited prion diseases in mice.

| **Drug Treatment** | **Scrapie mouse survival index** | **FFI mouse survival index** | **fCJD mouse survival index** |
| --- | --- | --- | --- |
| **IND24** | 2.47<sup>‡</sup> | 0.91 | 0.88 |
| **Anle128b** | 2.13<sup>‡</sup> | 0.95 | 1.12 |
| **IND24 + Anle128b** | 2.40<sup>‡</sup> | 0.74<sup>*</sup> | 0.84 |
| **Cellulose ether** | 3.24<sup>†</sup> | 0.97 | 1.00 |
| **PSCMA** | N/A | 0.92 | 1.07 |

Survival index = mean lifespan of treated mice/ mean lifespan of untreated controls. <sup>‡</sup>From
Burke *et al.*(11). <sup>†</sup>From Teruya *et al.*(12). <sup>\*</sup>P = 0.0017.

Untreated kiBVI^D178N^ and kiBVI^E200K^ mice spontaneously died of prion disease at ∼520 days and ∼650 days of age, respectively **(Figures 2a and 2b, black triangles)**. Paradoxically, kiBVI^D178N^ mice treated with a combination of Anle138b + IND24 developed disease more quickly (∼380 days) compared to untreated controls (∼520 days) (P = 0.0017). There was no statistically significant difference in the survival of any other treatment groups compared to untreated controls (**Figure 2** and **Table 1, columns 3-4**). Thus, none of the regimens tested provided significant therapeutic benefit in either FFI or kiBVI^E200K^ mice, despite displaying strong efficacy in mice infected with wild-type prions **(Table 1, compare column 2 *vs.* columns 3-4)** or inhibitory activity in mutant recombinant sPMCA reactions **(Figure 1)**.

**Figure 2:**
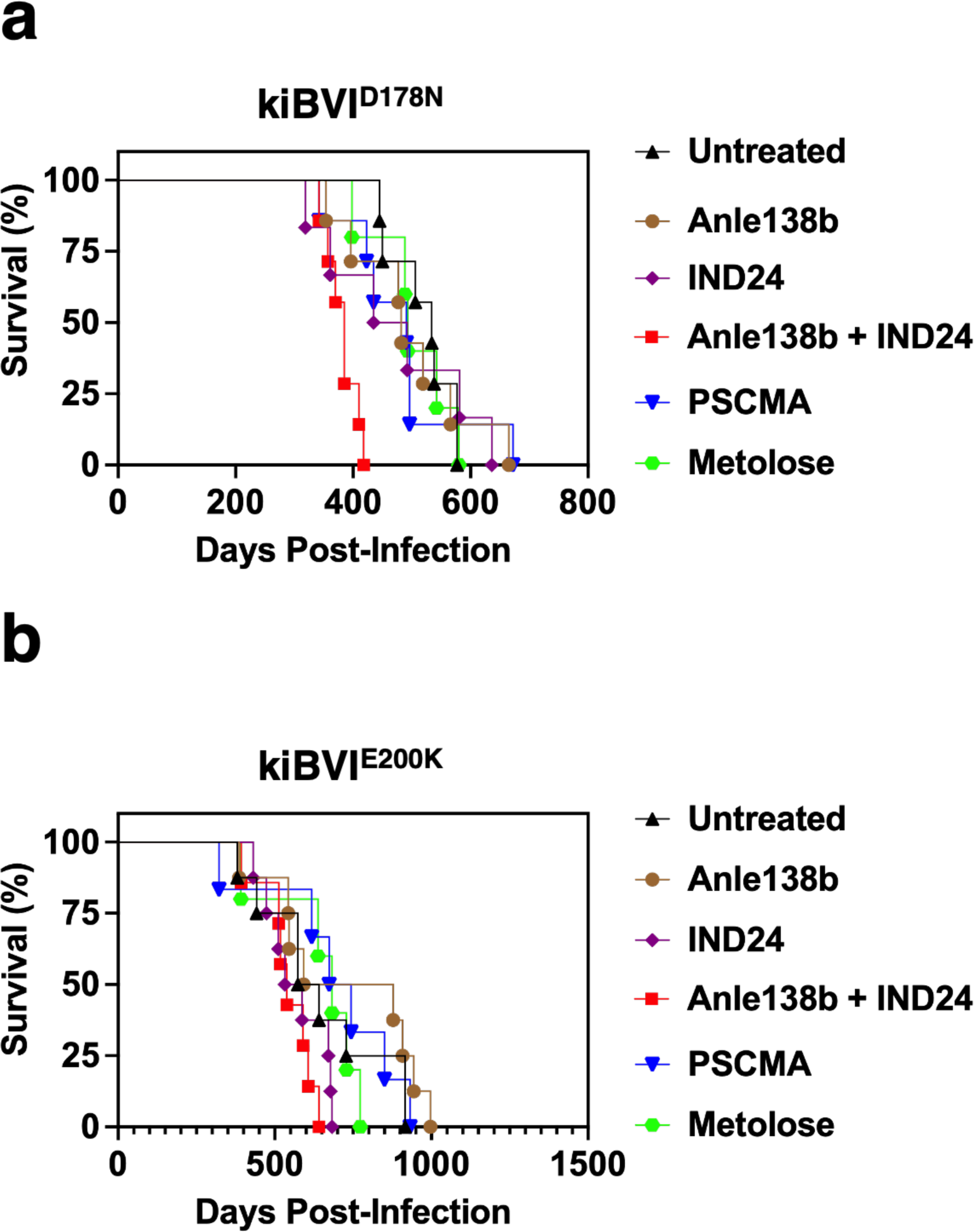
Effect of various drug treatments on survival in kiBVI^D178N^ and kiBVI^E200K^ mice. Survival curves of knock-in mice treated with various anti-prion drug regimens, as specified in the legend. **(A)** Survival curves of kiBVI^D178N^, and **(B)** kiBVI^E200K^ mice .

To investigate how the combination of Anle138b + IND24 might have decreased survival in kiBVI^D178N^ mice, we tested the effect of these two compounds on the kinetics of amyloid formation in unseeded QuIC assays with mutant recombinant BV PrP substrates. The results show that the combination of Anle138b + IND24 significantly accelerated D178N BV PrP amyloid formation compared to untreated reactions **(Table 2)** (P = 0.0495). These *in vitro* data suggest that the combination drug regimen may directly promote the process of D178N PrP^Sc^ formation.

**Table 2.**
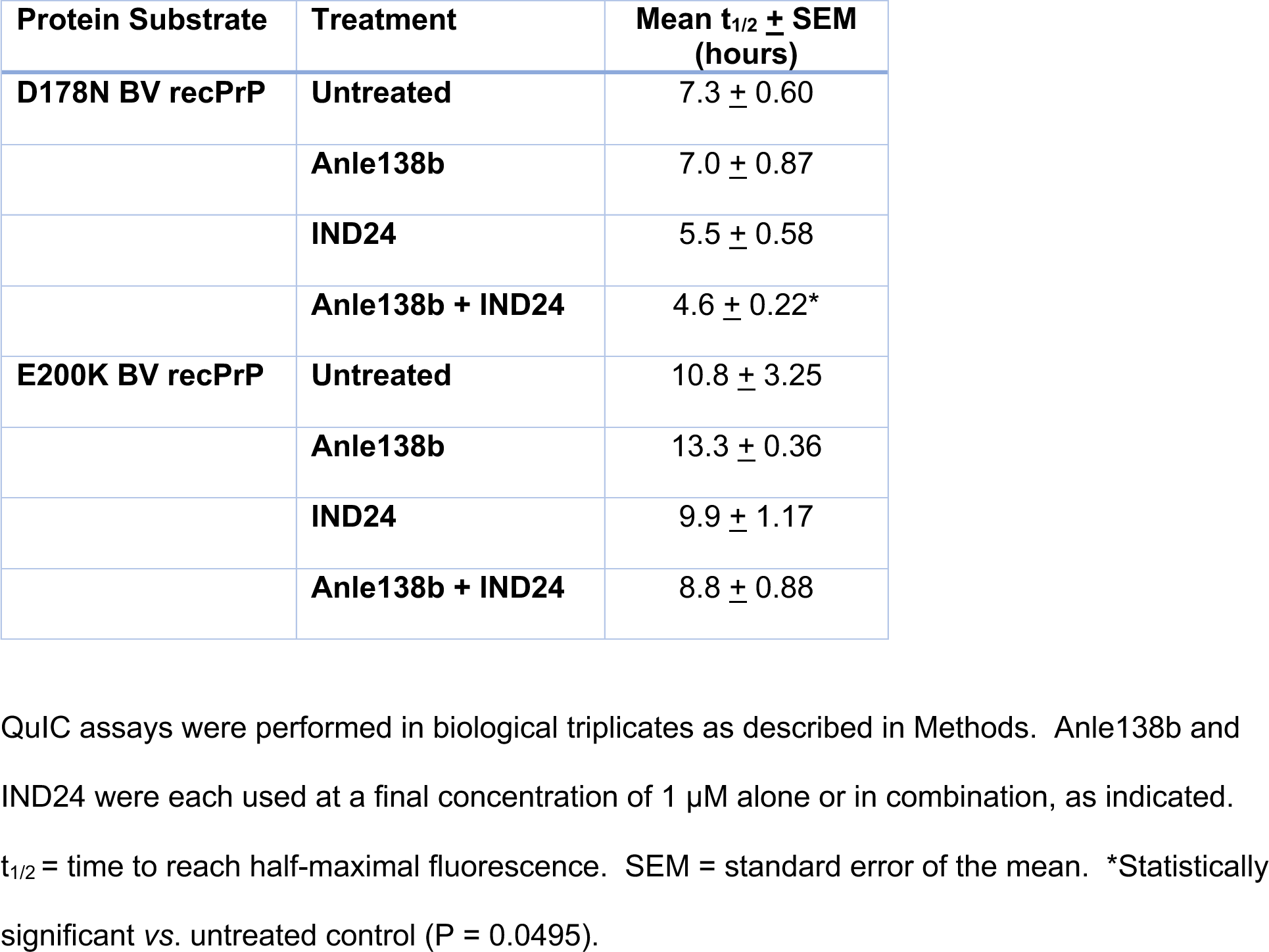
Effect of Anle138b and IND24 on QuIC assays with mutant PrP substrates.

| **Protein Substrate** | **Treatment** | **Mean $t_{1/2} \pm$ SEM (hours)** |
| --- | --- | --- |
| **D178N BV recPrP** | **Untreated** | $7.3 \pm 0.60$ |
| | **Anle138b** | $7.0 \pm 0.87$ |
| | **IND24** | $5.5 \pm 0.58$ |
| | **Anle138b + IND24** | $4.6 \pm 0.22^*$ |
| **E200K BV recPrP** | **Untreated** | $10.8 \pm 3.25$ |
| | **Anle138b** | $13.3 \pm 0.36$ |
| | **IND24** | $9.9 \pm 1.17$ |
| | **Anle138b + IND24** | $8.8 \pm 0.88$ |

QuIC assays were performed in biological triplicates as described in Methods. Anle138b and IND24 were each used at a final concentration of 1 μM alone or in combination, as indicated.
$t_{1/2}$ = time to reach half-maximal fluorescence. SEM = standard error of the mean. \*Statistically significant vs. untreated control (P = 0.0495).

Exposure to anti-prion drug regimens, including several of the regimens tested here, can lead to the selection and emergence of wild-type prions with altered biochemical and neuropathological strain properties (10, 11). To examine whether anti-prion drug therapy can also change the strain properties of mutant prions, we compared the migration patterns of protease-resistant PrP^Sc^ molecules as well as regional distributions of vacuolation in the brains of kiBVI^D178N^ and kiBVI^E200K^ mice in our control *vs.* treatment groups.

To detect protease-resistant PrP^Sc^ molecules in the brains of mutant mice, we treated brain homogenates of kiBVI^D178N^ and kiBVI^E200K^ mice from all the treatment groups with thermolysin (TL) and performed western blots with two different anti-PrP monoclonal antibodies, 27-33 (epitope = residues 142-149 mouse numbering) and EP1802Y (epitope = residues 217-226). We only detected TL-resistant PrP^Sc^ bands in a subset of knock-in mouse that developed spontaneous disease. In general, the majority of kiBVI^D178N^ mice had TL-resistant PrP^Sc^ in their brains, whereas few kiBVI^E200K^ mice did **(Figure 3 and Supplemental Figures S1-S6)**. When we compared the electrophoretic mobility of TL-resistant PrP^Sc^ bands from kiBVI^D178N^ mice in different treatment groups, we observed several differences in estern blots probed with mAb 27-33 **(Figure 3A, top blot)**. Most strikingly, whereas a ∼14 kDa TL-resistant band is present in untreated FFI mouse brains, IND24- and PSCMA-treated FFI mouse brains contain a ∼16 kDa TL-resistant band instead **(Figure 3A, top blot)**. In contrast, TL-resistant PrP^Sc^ bands showed similar electrophoretic mobility between individual kiBVI^D178N^ mice within each treatment group **(Supplemental Figures S1-S6)**. These results suggest that treatment with either IND24 or PSCMA induced the emergence of an alternative PrP^Sc^ conformation in kiBVI^D178N^ mice .

**Figure 3:**
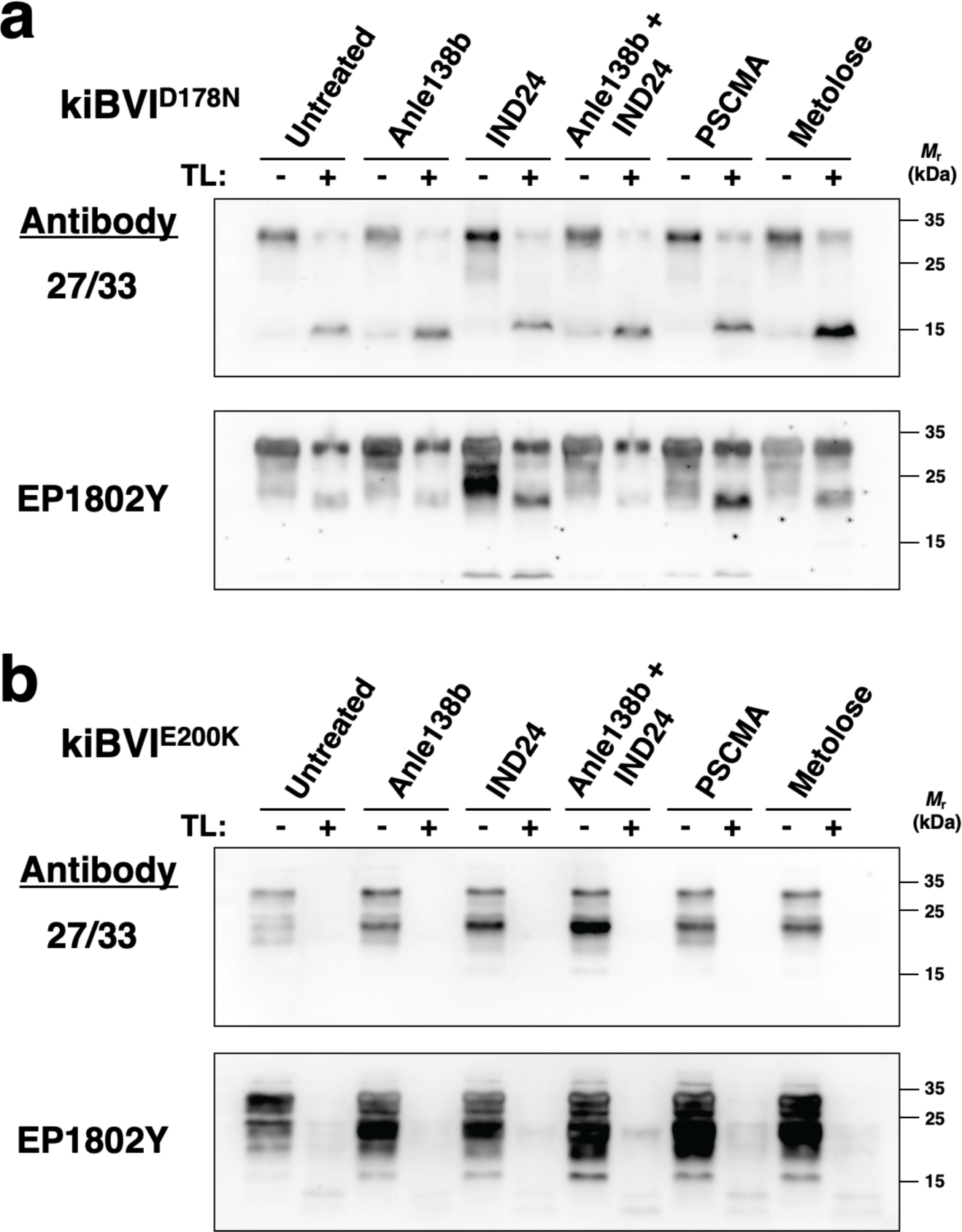
Mutant PrP^Sc^ molecules in brains of kiBVI^D178N^ and kiBVI^E200K^ mice treated with various anti-prion drug regimens. Western blot showing insoluble and thermolysin-resistant PrP^Sc^ molecules in brain homogenates of **(A) kiBVI^D178N^** and **(B)** kiBVI^E200K^ mice treated with various drug regimens, as indicated. Samples were either treated with thermolysin (TL) or water, as indicated. All samples were centrifuged to collect insoluble PrP. Within each panel, the top blot is probed with anti-PrP mAb 27-33 (epitope = residues 142-149 mouse numbering) and the lower blot is probed with mAb EP1802Y (epitope = residues 217-226).

We also observed differences in vacuolation profiles between various treatment groups and untreated controls in both kiBVI^D178N^ and kiBVI^E200K^ mice **(Figure 4)**. Specifically, kiBVI^D178N^ mice treated with either Anle138b + IND24 or PSCMA displayed different vacuolation profiles than untreated kiBVI^D178N^ mice, and kiBVI^E200K^ mice treated with IND24, Anle138b + IND24, PSCMA, or Metolose displayed different vacuolation profiles than untreated kiBVI^E200K^ mice **(Table 3)**. A particularly striking example of drug-induced change in neuropathology was observed in the corpus callosum of kiBVI^E200K^ mice . Whereas untreated kiBVI^E200K^ mice displayed abundant vacuolation in the corpus callosum, kiBVI^E200K^ mice treated with either IND24 alone or Anle138b + IND24 showed little to no vacuolation in the corpus callosum **(Figure 5)**. Taken together, the biochemical and neuropathological strain typing assays provide evidence that all the drug regimens used in this study except Anle138b monotherapy altered the strain properties of mutant prions in both kiBVI^D178N^ and kiBVI^E200K^ mice.

**Figure 4:**
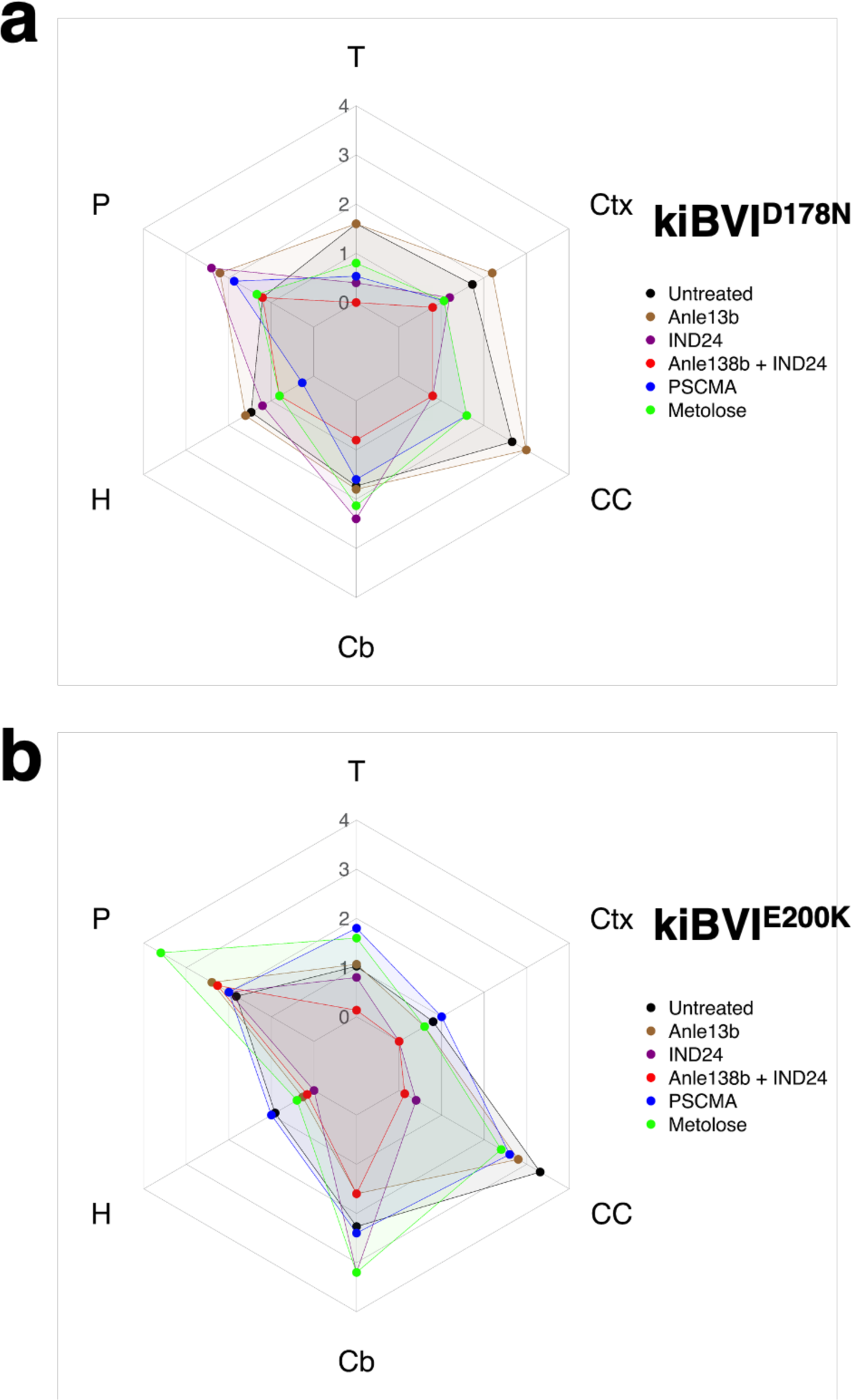
Regional vacuolation profiles in brains of kiBVI^D178N^ and kiBVI^E200K^ mice treated with various anti-prion drug regimens. Mean vacuolation scores (n = 2-7) in various brain regions of mice treated with various drug regimens, as specified in the legends. T = Thalamus, Ctx = cerebral cortex, CC = corpus callosum, Cb = cerebellum, H = hippocampus, P = pons. **(A)** Regional vacuolation profiles of kiBVI^D178N^ mice, and **(B)** regional vacuolation profiles of kiBVI^E200K^ mice .

**Figure 5:**
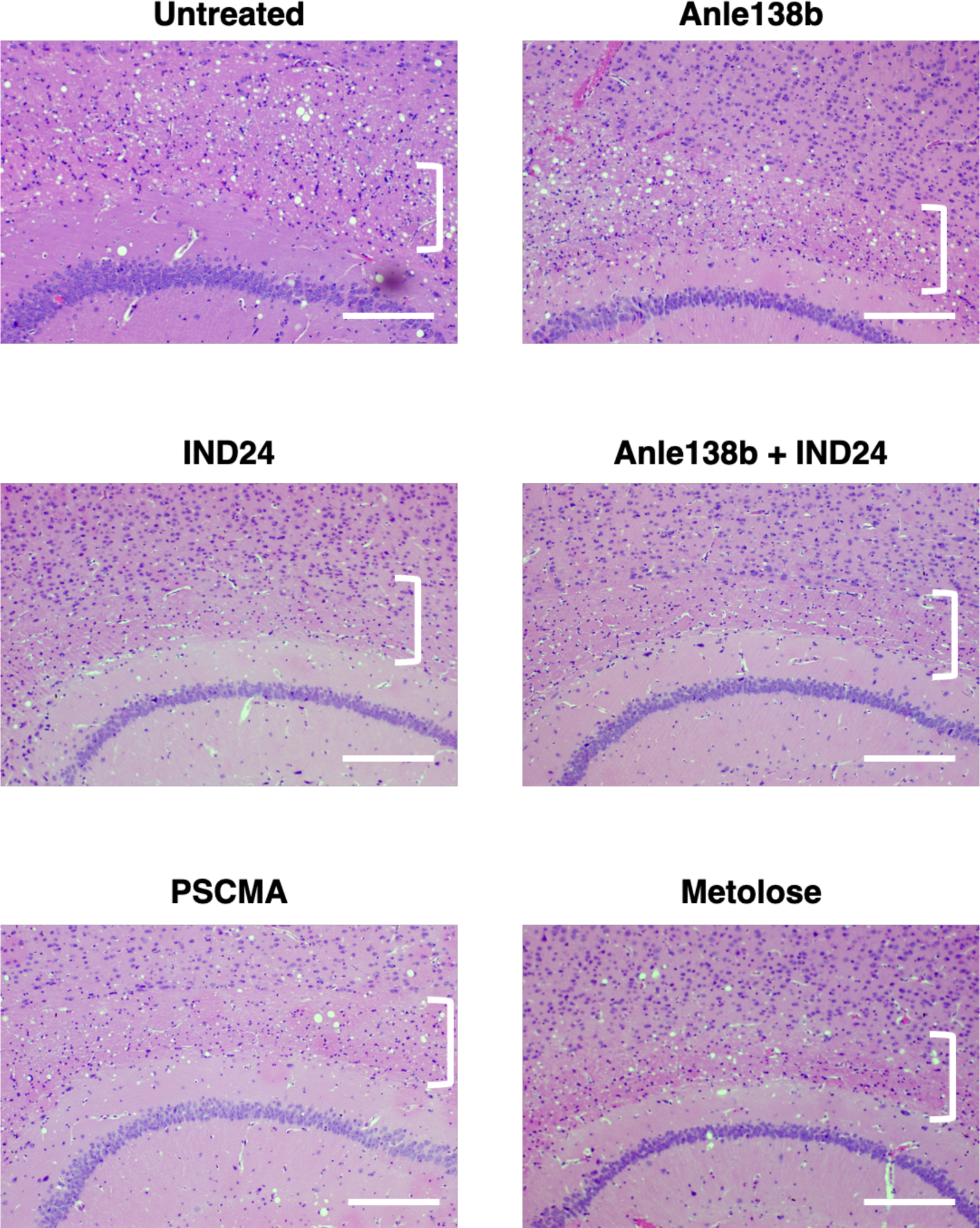
Neuropathology of corpus callosum in kiBVI^E200K^ mice treated with various anti-prion drug regimens. Representative microscopic images of brain sections of kiBVI^E200K^ mice treated with various drug regimens, as specified, stained with hematoxylin and eosin. Square brackets indicate the location of the corpus callosum (in between the cerebral cortex and hippocampus). Horizontal scale bar = 200 μm.

**Table 3:** P-values of significant differences in vacuolation scores of treatment groups *versus* untreated controls in different brain regions.

|  | T | Ctx | CC | Cb | H | P |
| --- | --- | --- | --- | --- | --- | --- |
| **kiBVI<sup>D178N</sup> mice** |  |  |  |  |  |  |
| Anle138b |  |  |  |  |  |  |
| IND24 |  |  |  |  |  |  |
| Anle138b + IND24 | 0.038 |  |  | 0.021 |  |  |
| PSCMA |  |  |  |  | 0.041 |  |
| Metolose |  |  |  |  |  |  |
| **kiBVI<sup>E200K</sup> mice** |  |  |  |  |  |  |
| Anle138b |  |  |  |  |  |  |
| IND24 |  |  | 0.024 |  |  |  |
| Anle138b + IND24 | 0.004 | 0.013 | 0.001 |  | 0.028 |  |
| PSCMA | 0.022 |  |  |  |  |  |
| Metolose |  |  |  |  |  | 0.012 |

Brain regions displaying statistically significant differences in vacuolation scores between treated mice vs. untreated controls are highlighted with p-values (n = 2-7 per group). T = Thalamus, Ctx = cerebral cortex, CC = corpus callosum, Cb = cerebellum, H = hippocampus, P = pons. Where not shown, P > 0.05.

## Discussion

Here we report the first drug trial in knock-in animal models of FFI and fCJD with shortened lifespans. We tested four drug regimens that have previously been shown to inhibit the accumulation of wild-type PrP^Sc^ and significantly prolong incubation time in scrapie-infected animals(10–12, 34) and one compound that potently inhibits mutant PrP^Sc^ propagation in recombinant sPMCA reactions. Surprisingly, we observed that none of these drugs inhibited the accumulation of mutant PrP^Sc^ in or significantly extended the lifespan of either FFI or kiBVI^E200K^ mice (Anle138b and PSCMA increased lifespan modestly in fCJD). Notably, all the compounds we tested belong to different chemical classes, cross the blood-brain barrier, and are well-tolerated by mice. Three of the compounds tested (IND24, Anle138b, and Metolose cellulose ether) represent the most efficacious prophylactic drugs currently known for wild-type scrapie prions, extending scrapie incubation time between 2-to 4-fold(9–12, 34). A fourth compound, PSCMA, has not yet been tested in prion-infected animals, but potently inhibits D177N propagation in recombinant sPMCA reactions *in vitro* and blocks PrP^C^-dependent Aβ oligomer toxicity *in vivo*(35).

It has been previously shown that non-specific clearance therapies (rapamycin and IVIG) but not Anle138b can delay the onset of disease in a transgenic mouse model of Gerstmann-Sträussler-Scheinker (GSS) syndrome(36–38). Our current study differs from these prior studies because (1) GSS is a phenotypically distinct inherited prion disease characterized by the accumulation of amyloid plaques not seen in FFI or fCJD, and (2) we used knock-in rather than overexpression animal models. Theoretically, using animal models with lower expression levels and fewer plaques should improve our ability to detect a therapeutic effect and therefore increase our confidence about the negative results observed in this study. A more recent study testing Anle138b in alternative knock-in mouse models of FFI and fCJD expressing mutant mouse PrP molecules rather than bank vole PrP molecules was limited by the relatively normal lifespan of those mutant mice(25).

It has been previously shown that drug-resistant wild-type prions with altered PrP^Sc^ conformation and strain characteristics can emerge in response to anti-prion drug therapy(10) (11). In this study, we observed that mutant prions can also undergo strain adaptation in response to drug therapy *in vivo*, based on changes in biochemical and neuropathological strain-typing assays. At the same time, we did not observe a survival benefit for any of the anti-prion therapies tested. Together, these results suggest that drug-resistant conformers may emerge more rapidly during the replication of mutant prions compared to wild-type prions. It is worth considering, as an alternative to the strain selection hypothesis, the possibility that each drug may preferentially inhibit prion replication in specific brain regions, and that lifespan is determined by disease progression in the regions not impacted by each drug. However, this alternative hypothesis would not explain the differences in mobility of TL-resistant PrP^Sc^ bands in IND24-treated and combination-treated mice vs. untreated mice.

An unexpected result in our study was that combination therapy with Anle138b and IND24 paradoxically accelerated disease in kiBVI^D178N^ and kiBVI^E200K^ mice. This effect is not due to drug toxicity because (1) this combination regimen does not accelerate disease in wild-type mice(11), and (2) knock-in mice with combination drug-accelerated disease have mutant PrP^Sc^ in their brains. We observed that the combination of Anle138b and IND24 accelerated mutant PrP aggregation in QuIC reactions, suggesting that the combination regimen may also increase the kinetics of mutant PrP^Sc^ formation *in vivo*. It is possible that strain adaptation of mutant prions induced by combination therapy selects for a conformer that is either more toxic or difficult to degrade in neurons. Regardless of mechanism, this result cautions that drug regimens which successfully inhibit wild-type prion replication may paradoxically harm patients with inherited forms of prion disease.

The major conclusion of our study is that drugs that effectively treat wild type prion disease are inactive against two different mutant prions in knock-in mouse models of inherited prion disease. One hypothesis for the ineffectiveness of the repurposed drugs is that the spontaneous formation of mutant prions and templated formation of wild-type prions may use different mechanisms. Mutant prions can form without seed or cofactor molecules, whereas cofactor molecules are required to form and propagate wild-type prions(24). Cofactor molecules are even required by wild-type PrP^C^ substrate molecules to propagate the conformation of mutant PrP^Sc^ seeds(24). Consistent with this hypothesis, the host translational response of mice expressing mutant prions is strikingly different from the translation response of mice infected with wild-type prions(39, 40). An alternative hypothesis is that none of the three repurposed drugs tested work on the mutant prion strains in kiBVI^D178N^ and kiBVI^E200K^ mice. Each of the three of these drugs has been previously shown to inhibit some prion strains but not others: IND24 inhibits RML, Me7, and chronic wasting disease (CWD), but not sCJD prions(10); Metolose inhibits 263K and CWD, but not RML prions(12, 41, 42); and Anle138b inhibits RML, but not 263K prions(43).

A critical lesson from this study is that treatments for inherited prion diseases should be identified and/or developed by specifically targeting mutant prions rather than repurposing drugs that work on wild-type prions. Developing valid cell-based assays to screen candidate compounds for their ability to inhibit mutant prions would greatly facilitate such efforts since biochemical assays may not accurately predict *in vivo* efficacy. We observed that PSCMA potently inhibited D177N PrP^Sc^ propagation in recombinant sPMCA reactions, but PSCMA had no therapeutic effect *in vivo*. Our results also suggest that the novel knock-in models of FFI and fCJD that we used for these studies may be generally advantageous for pre-clinical *in vivo* testing because they have physiological PrP expression levels and shortened lifespans, thereby providing an objective and quantifiable endpoint for evaluating therapeutic efficacy, as recently suggested by Vallabh and Minikel(25). Finally, our results suggest that less easily administered therapies which lower PrP levels (such as anti-sense oligonucleotides delivered by lumbar puncture) may ultimately be required to escape the ability of both wild-type and mutant prions to undergo strain-adaptation leading to anti-prion drug failure(32, 44, 45).

In summary, our findings show that several anti-prion drugs with activity against wild-type prions have no beneficial effect on disease-free survival of two different knock-in mouse models of inherited prion disease. Our work highlights the importance of developing valid drug screening assays and alternative therapies specifically for inherited prion disease.

