## Supplemental Figures for "Anti-prion drugs do not improve survival in knock-in models of inherited prion disease"

**Supplemental figures S1-S6: Mutant $PrP^{Sc}$ molecules in brains of** $kiBVI^{D178N}$ and $kiBVI^{E200K}$ mice**. Figure S1:** Untreated mice. **Figure S2:** mice treated with Anle138b. **Figure S3:** mice treated with IND24. **Figure S4:** mice treated with a combination regimen of Anle138b + IND24. **Figure S5:** Mice treated with PSCMA. **Figure S6:** Mice treated with Metolose. **All Figures:** Western blots showing insoluble and thermolysin-resistant PrPSc molecules in brain homogenates of knock-in $kiBVI^{D178N}$ and $kiBVI^{E200K}$ mice treated with various drug regimens, as indicated. Samples were either treated with thermolysin (TL) or water, as indicated. All samples were centrifuged to collect insoluble PrP. In each panel, the top blot was probed with anti-PrP mAb 27-33 and the bottom blot was probed with mAb EP1802Y. **(A)** Blots of D178N PrP molecules in $kiBVI^{D178N}$ mice , **(B)** blots of E200K PrP molecules in $kiBVI^{E200K}$ mice .

### Supplemental Figure S1:  Untreated

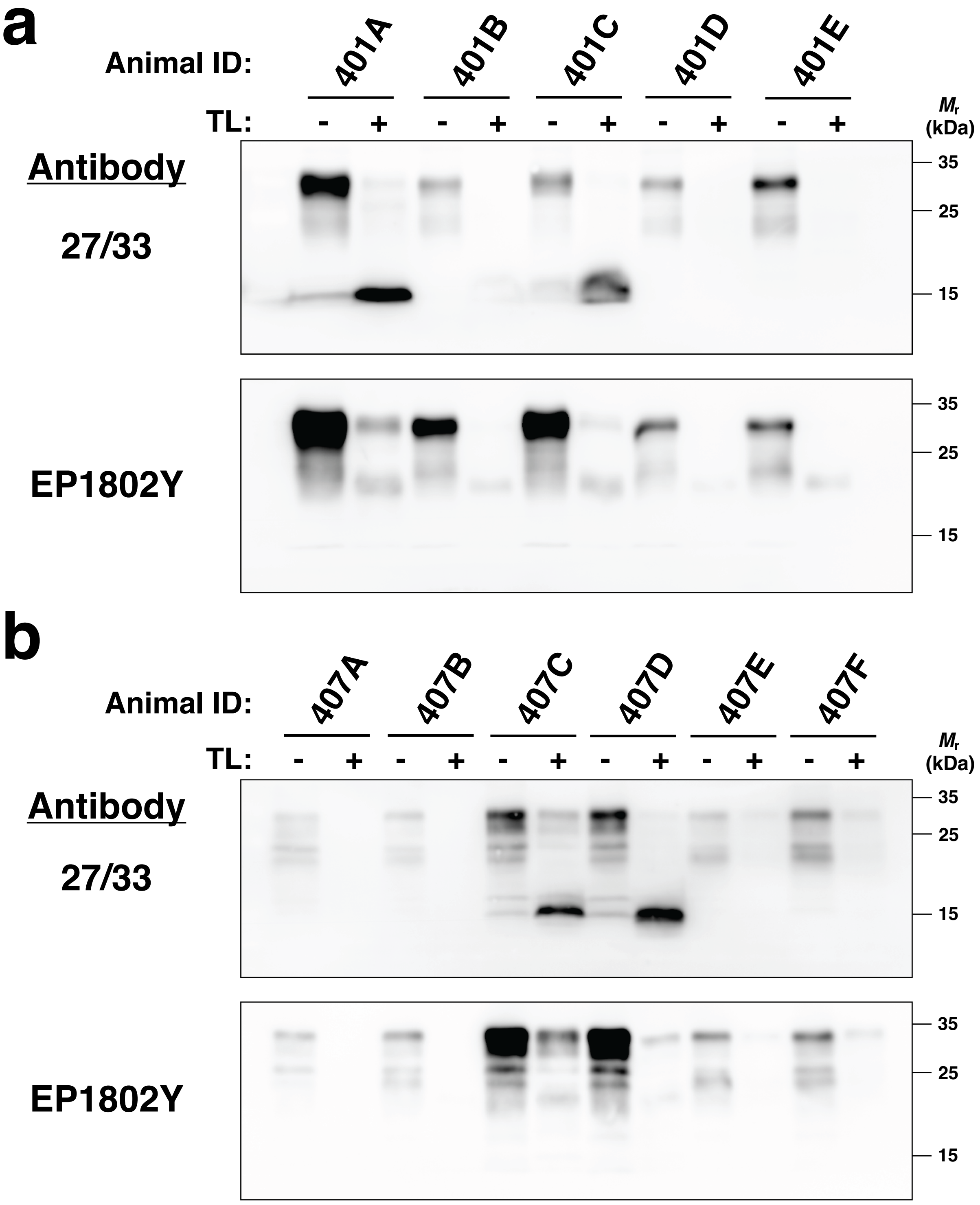

### Supplemental Figure S2: Anle 138b

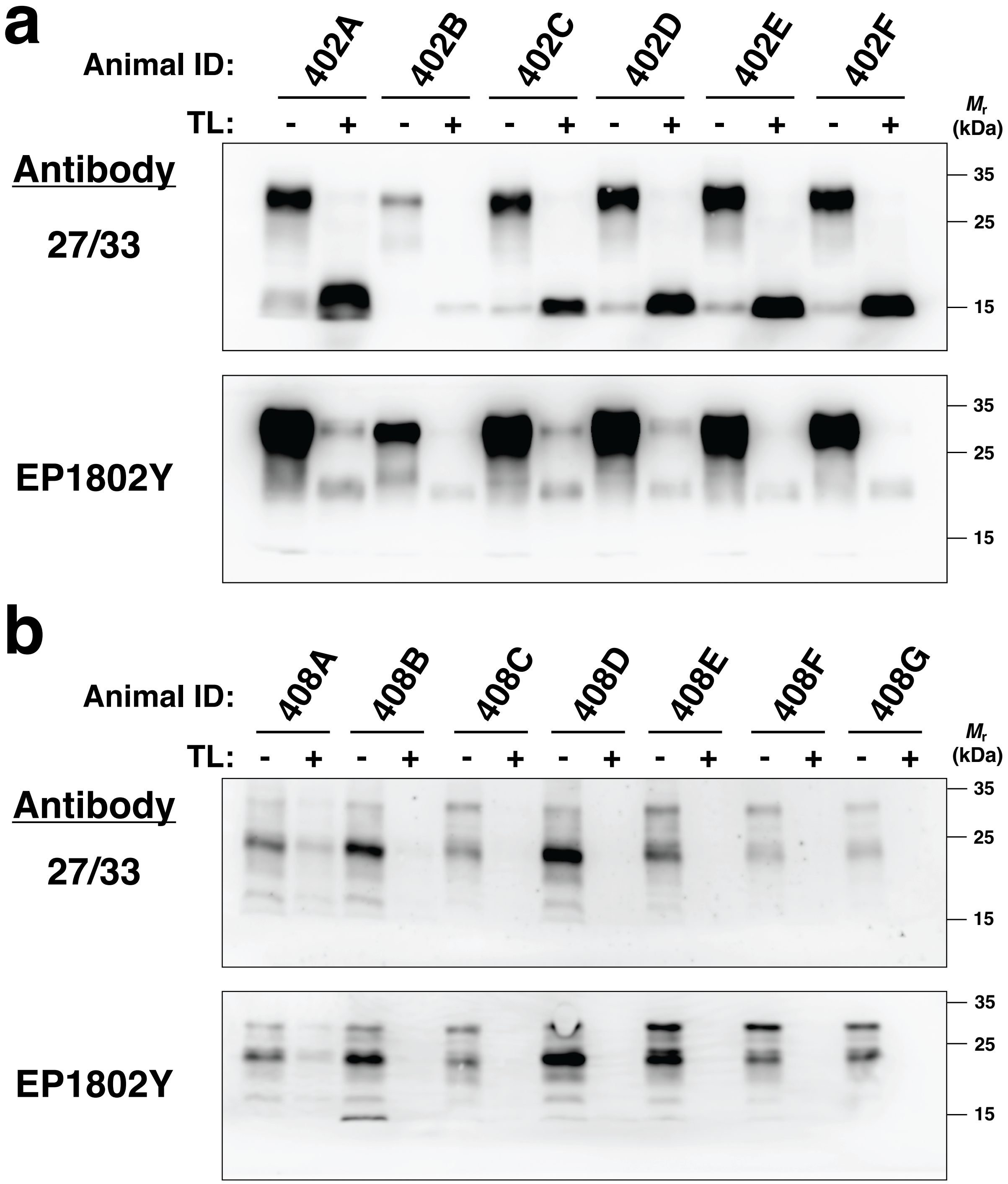

### Supplemental Figure S3: IND24

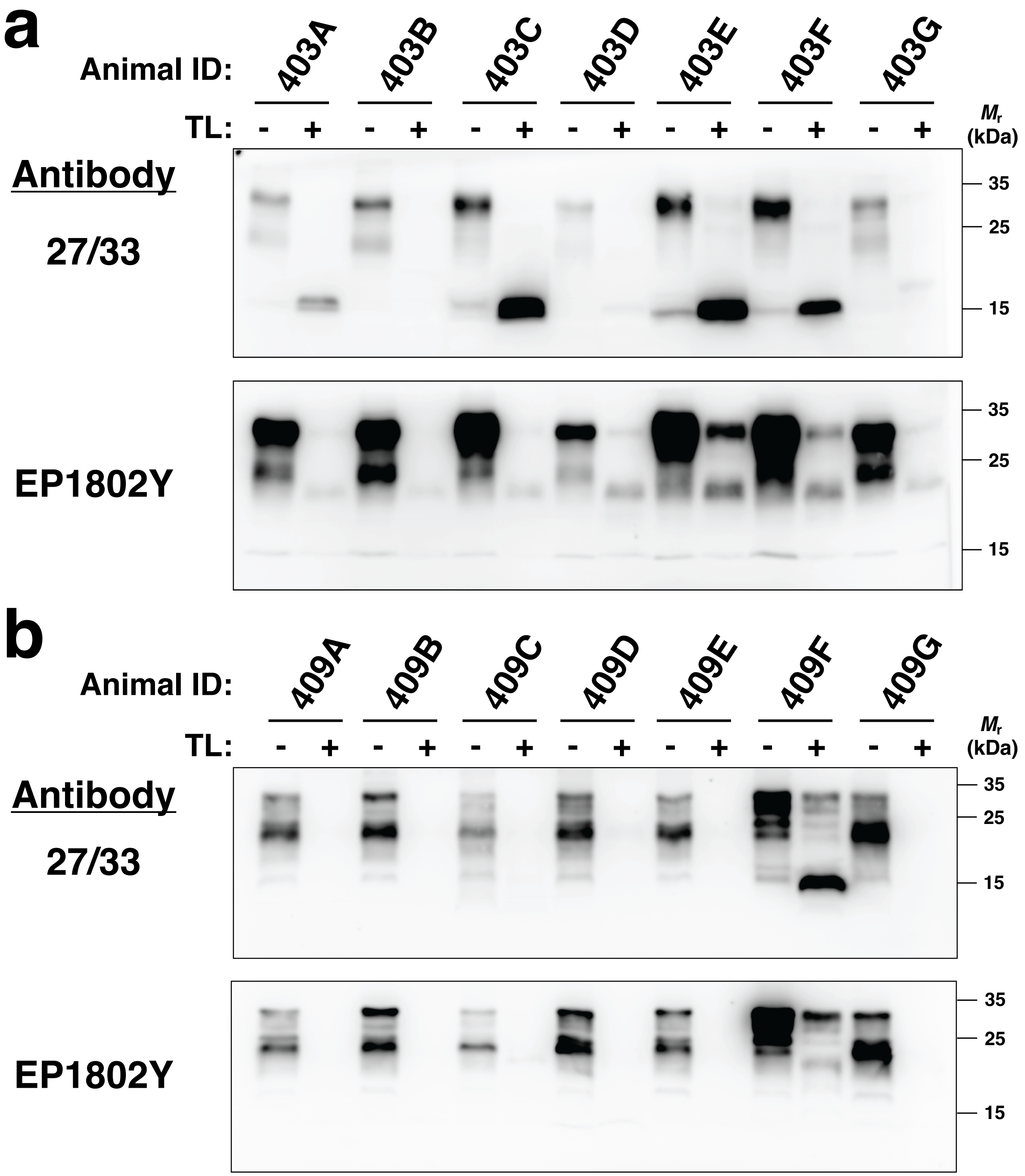

#### Supplemental Figure S4: Anle138b + IND24

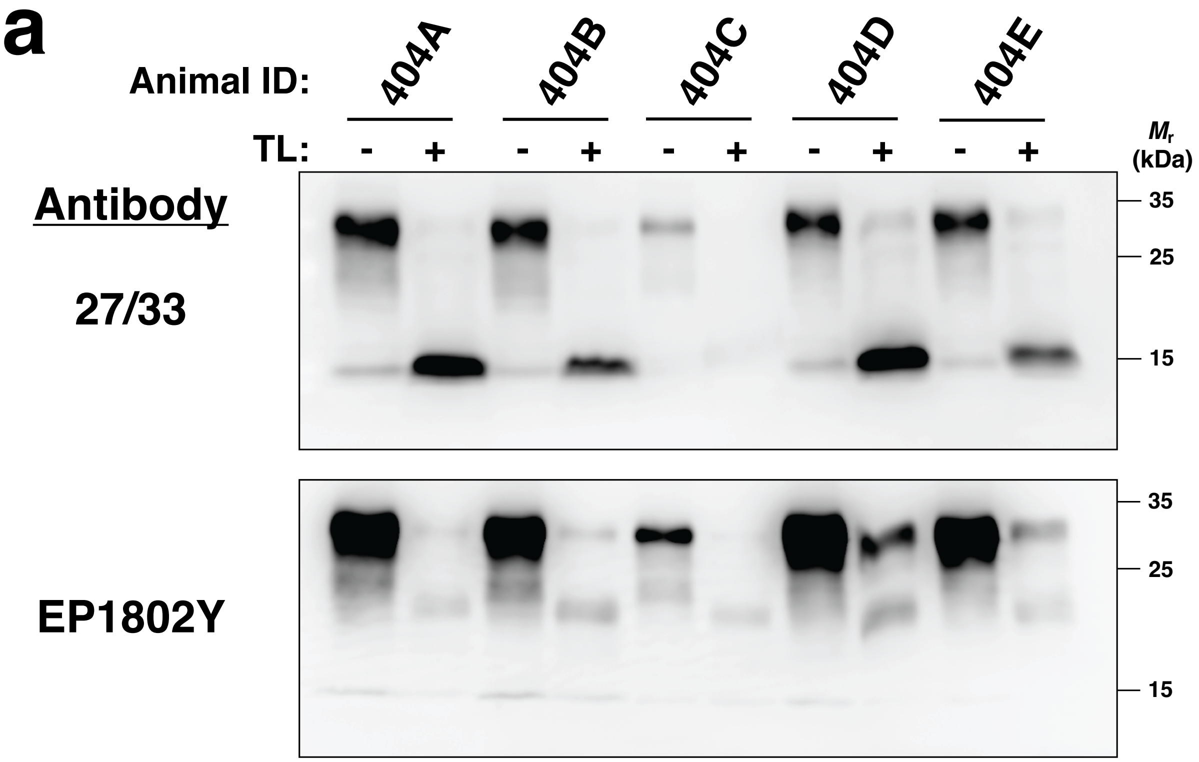


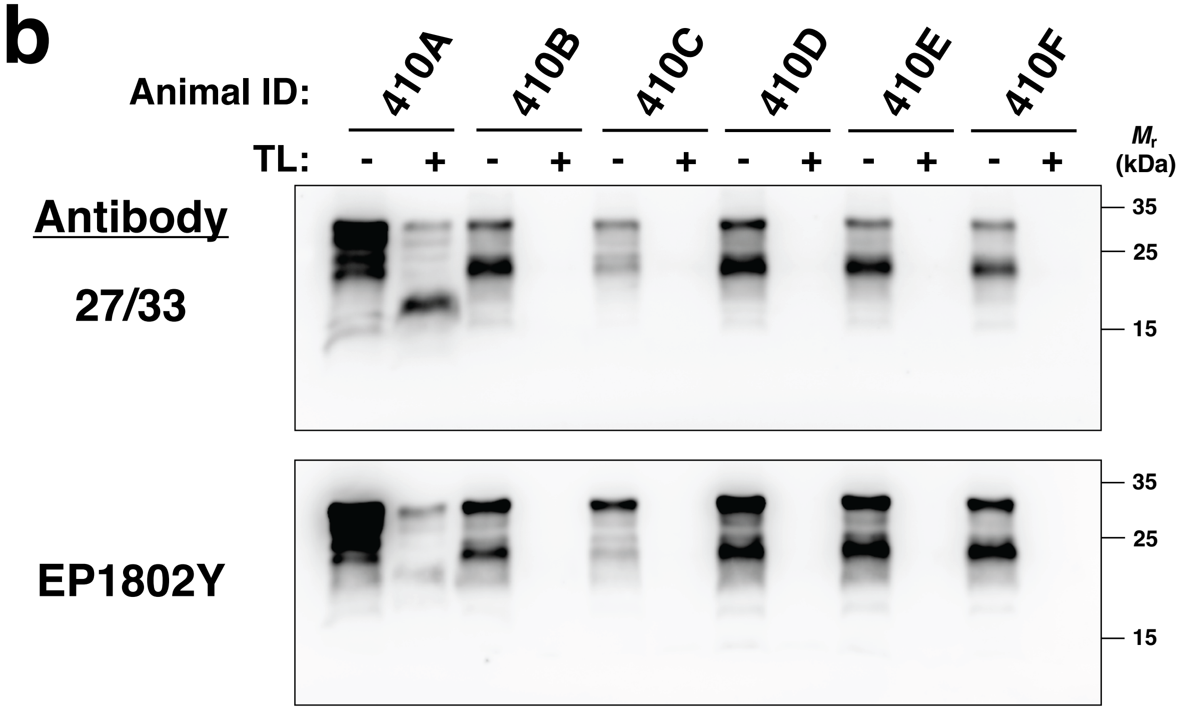

### Supplemental Figure S5:  PSCMA

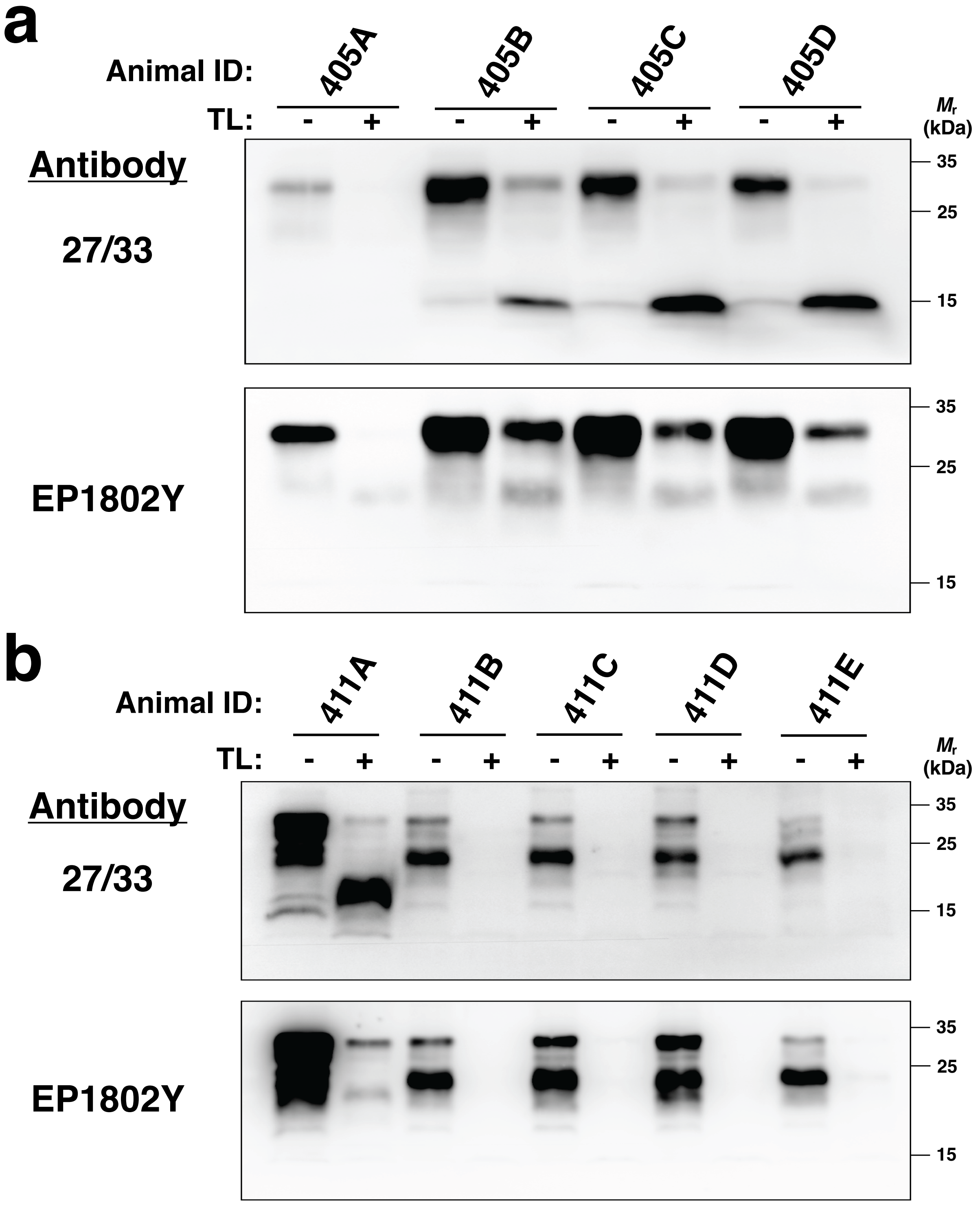

#### Supplemental Figure S6: Metolose

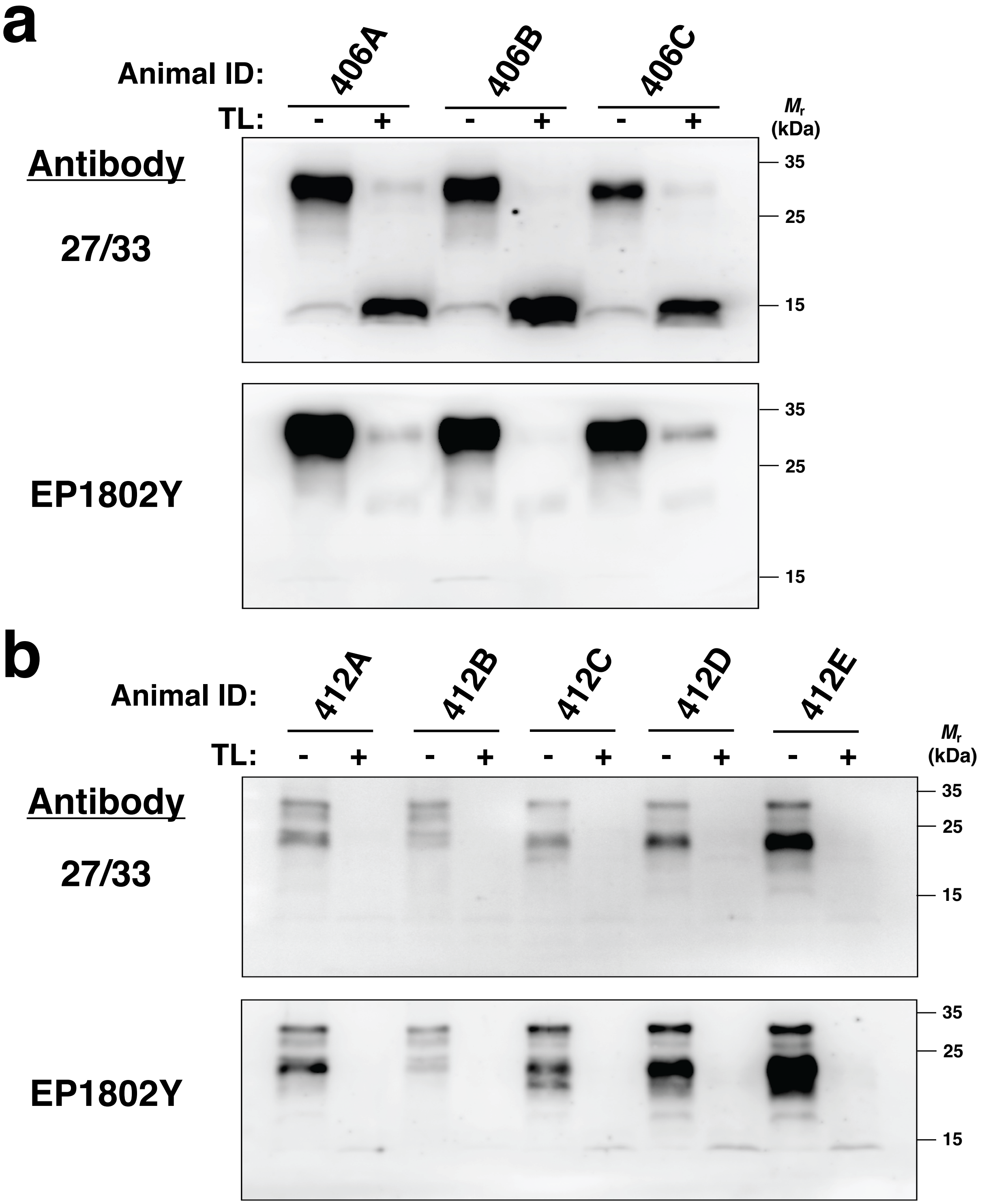
